# An Open Quantum System Approach to Bicoid Gradient Dynamics

**DOI:** 10.1101/2024.09.20.614139

**Authors:** Irfan Lone

## Abstract

A prototypical morphogen gradient that plays a key role in the early embryonic development of fruit flies, by providing positional information to cells, is that of the transcription factor Bicoid (Bcd). Recently a one-dimensional quantum walk model has been utilised to explain its multiple dynamic modes observed through fluorescence correlation spectroscopy (FCS) studies using a closed quantum system approach. In this work we use an open quantum system approach to the dynamics of the Bcd gradient formation and show that exactly the same dynamics are obtained through this more rigorous analysis. We then use the thus obtained expression for the fast dynamic modes to explain the Bcd transcription factor search times for binding to the promoter regions along the DNA. Specifically, we find that the large values of diffusivity allowed by quantum mechanics can avoid the paradox of faster-than-diffusion association rates without any need for the transcription factor to constantly alternate between 1*D* and 3*D* diffusion-based search processes. This might help explain the fast and precise transcriptional response elicited by such factors. We conclude that, since many transcription factors share a common search strategy for target gene regulatory regions, our mechanism may have a wide range of applicability.

---

Morphogen gradients play a crucial role in the early embryonic development of multicellular organisms by providing positional information to cells [1–10]. A prototypical morphogen gradient is that of the transcription factor Bicoid (Bcd), formed in the early Drosophila melanogaster embryo and plays a key role in the determination of embryonal axis of the organism [11]. Although the Bcd gradient has been studied for a very long time [11–15], the biophysical mechanisms behind its establishment are still not completely understood. In 1970, Francis Crick suggested that morphogen gradients could be a consequence of diffusion [16], frequently assumed to be the mechanism associated with the biological transport processes. Crick introduced a model in which there is a constant production of morphogen at one end and its spatially uniform degradation throughout the system setting up a concentration gradient in three-dimensional space [16]. The state of such a system can be described as a classical random walk process, involving both diffusion and degradation, through the use of following reaction–diffusion equation,

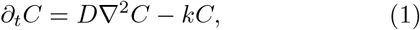

where *D* and *k* are, respectively, the diffusion and degradation constant of the morphogen, and *C* denotes its concentration. Further extensions of Crick’s ideas have essentially taken two directions. On the one hand the discovery of the Bcd gradient in the early fruit fly embryo [11] has motivated several diffusion-based models [17]. On the other hand, fluorescence and single-molecule imaging based experiments [18–20] have inspired theories using explicit kinetic Markovian schemes based on the chemical master equation [21]. Further extensions accounting for both intrinsic noise [22–26] and diffusion [27–34] build upon Crick’s work. Regardless of the specific model details, all of these models share the common diffusion-degradation paradigm of Crick’s model in which the time evolution of the concentration fields, or probability densities, for the system is given by a deterministic reaction-diffusion equation or some coarse-grained approach based on a master equation [21].

In contrast, a recent experimental observation, utilising fluorescence correlation spectroscopy (FCS) and perturbative techniques, has revealed multiple modes of Bcd transport at different spatial and temporal locations across the embryo [35]. Exploring the different dynamic modes for Bcd it has been observed that in the cytoplasm the slow dynamic mode is similar across the embryo while as the fast one shows an increased diffusivity from the anterior to the posterior side [35]. Similarly, in the nuclei the Bcd diffusivity shows a significant, though comparatively small, increase in the fast dynamic mode from the anterior to the posterior regions, with the slow component remaining essentially unchanged [35]. These observations, and a few others, have recently been explained through a quantum-classical treatment of Crick’s model [36]. In the quantum-classical treatment, the degradation of Bcd is modelled as a unitary noise that is intrinsic and that does not cause any entanglement with the environment, so that the system remains in an essentially pure state during the course of its evolution. The degradation thus acts like a classical fluctuating field such that the dynamics of the system is unitary, yet stochastic [37].

Thus, the state of the system at any given time in such a model is represented by a vector Ψ in a finitedimensional Hilbert space [38]. Molecules are represented, just like in particle-based reaction–diffusion schemes *ℌ* [39], by point particles undergoing a Markovian quantum diffusion process in presence of a unitary noise that introduces a degree of stochasticity into their dynamics. One can therefore borrow the well-known results for diffusion dynamics from the theory of quantum Markov processes [40]. Thus, the model is constructed in a bottom-up way based on the theory of quantum Markov processes and analytically solved for a one-dimensional case [36]. It has therefore been shown that the multiple dynamic modes of the Bcd gradient are a consequence of a quantum-classical dynamics in which transient quantum coherences play an essential role. Such a treatment of the Bcd gradient has been carried out by assuming it to be a closed quantum system with a pure state Ψ so that its density operator is given by

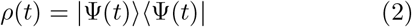

In this Letter we shall use an open quantum system approach [40] to the problem of Bcd gradient formation and demonstrate that the final expression for the dynamic modes is comparable to that obtained through the closed quantum system hypothesis. We shall then use the derived expression for diffusivity to explain the Bcd transcription factor search times for binding to the promoter regions along the DNA.

Motivated in part by the experimental observation of multiple dynamics modes accompanying the process of Bcd gradient formation [35], a quantum walk model of Bcd gradient formation [36] is proposed in the following manner (Fig. 1). Compared to the usual classical random walk models, the coin toss is replaced by a chirality degree of freedom which can take two values denoted |+⟩ and |−⟩, for rightand left-handed chirality, respectively [41]. A Bcd molecule of chirality state |+⟩ can move one step to the right at a time while that of chirality |−⟩ can move to the left along the lattice or they can also get degraded, leading to the setting up of a concentration gradient in one-dimensional space [42].

**FIG. 1.**
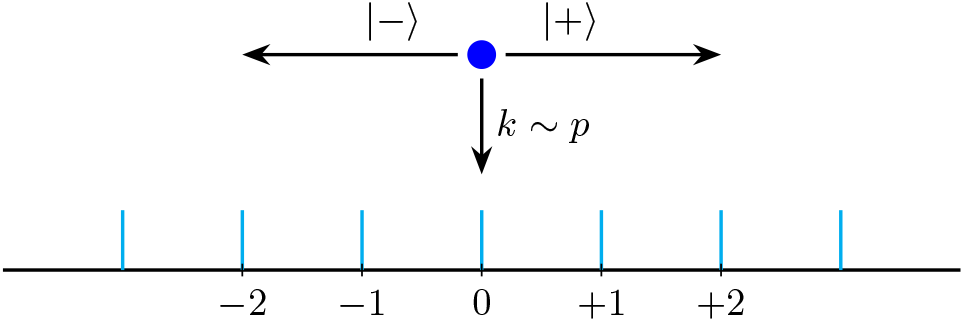
Schematic of the quantum-walk model. A Bcd molecule (shown by blue colored dot) can, respectively, make a transition to the right or to the left depending on its chirality state of |+⟩ or |−⟩ or it can get degraded at a rate *κ*.

The degradation of Bcd molecules is modelled by means of a unitary noise (Fig. 2) such that (i) When a Bcd molecule resides only briefly at a given site, it may not get internalized and degraded but instead survive and make a transition to the neighboring sites, |+⟩ to right and |−⟩ to the left. (ii) When a particle of chirality |+⟩ is internalized at a given site, e.g. *n* = 0, then there is no probability flux towards the nearby site, *n* = +1. Since the upper component of spinor at *n* = 0 sends probability flux towards *n* = −1, in order to conserve flux, the outgoing probability flux from the lower component at *n* = 0 must be diverted to the upper component at the same site. Thus the transition that should have taken place is turned into a self-loop for that step of the walk. However, there is still a flow of probability flux towards the site *n* = − 1 due to the particles of |−⟩ chirality state. (iii)When a particle of chirality |−⟩ is internalized at a given site, *n* = 0 again, then there is no flow of probability flux towards the nearby site on the left, *n* = −1, and the transition that should have taken place is again turned into a self-loop for that step of the walk. However, there is still a flow of probability flux towards the site *n* = +1 due to the particles of |+⟩ chirality state (iv) When particles of both chirality states reside for appreciably longer periods of time at a given site, they may both get internalized and degraded. In such a situation there is no flow of probability flux in either direction. In this unitary noise model, we denote the probability per time step of the Bcd molecule of a given chirality getting degraded by *p*. Thus, the parameter *p* quantifies noise in our model of the Bcd system. One can thus state that *p*^2^ represents the probability that Bcd molecules don’t make a transition in either direction, (1 −*p*)^2^ the probability that Bcd molecules make a transition in both directions and *p*(1 −*p*) the probability that Bcd molecules make a transition in one of the two possible directions [36].

**FIG. 2.**
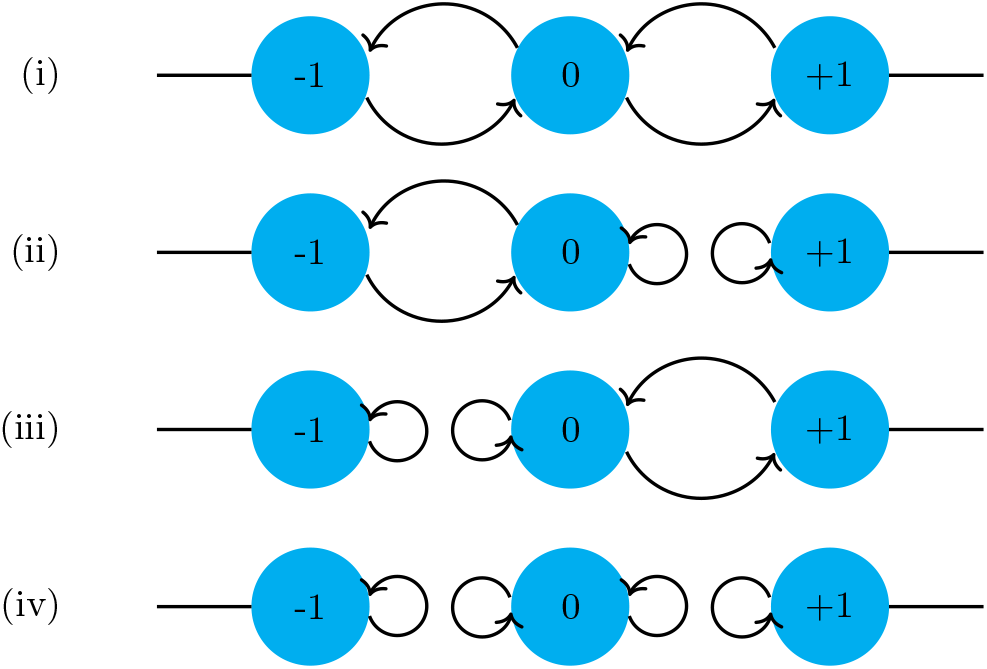
Schematic of the unitary noise in the Bcd system. The curved arrows depict the flow of probability flux between sites. Lattice sites, labelled by *n* = 0, ±1, ± 2, …, correspond to the positions (states) of the Bcd particles in motion. (i) When a Bcd molecule resides only briefly at a given site, it may not get internalized and degraded but instead survive and make a transition to the neighboring sites. (ii) When a particle of chirality |+⟩ is internalized at a given site, e.g. *n* = 0, then there is no probability flux towards the nearby site, *n* = +1. (iii) When a particle of chirality |−⟩ is internalized at a given site, *n* = 0 again, then there is no flow of probability flux towards the nearby site on the left, *n* =− 1. (iv) When particles of both chirality states reside for appreciably longer periods of time at a given site, they may both get internalized and degraded. In such a situation there is no flow of probability flux in either direction.

Let ℌ_*n*_ be the position Hilbert space of the system. In our model ℌ_*n*_ has a support on the space of eigenfunctions |*n*⟩ corresponding to the sites *n* ∈ ℤ on the lattice, ℤ being a set of integers. This position Hilbert space ℌ_*n*_ is augmented by a ‘coin’ space ℌ_*c*_ spanned by the two basis states {|+⟩, |−⟩}. The states of the total system are thus in the Hilbert space ℌ_*s*_ = ℌ_*n*_ ⊗ *ℌ*_*c*_. Let ℌ_*e*_ denote the Hilbert space of the environment spanned by the basis 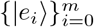, where *m* is the dimension of the environment Hilbert space [43]. The total Hilbert space of the problem is thus given by

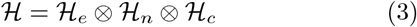

The state of the system is then obtained by tracing out over the environmental degrees of freedom as

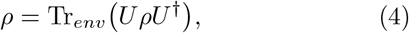

rather than from Eq. (2). The global unitary *U* acts both on the system and the environment Hilbert spaces. Now one may assume that the state of the whole system is initially given by

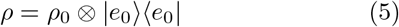

Then one can write

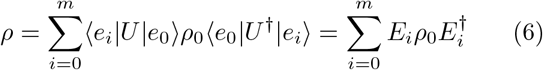

where *E*_*i*_ = ⟨*e*_*i*_ *U e*_0_ ⟩, *i* = 0, 1, …, *m*, are the so-called Kraus operators [44]. The Kraus operators follow the completeness relation, which arises from the requirement that

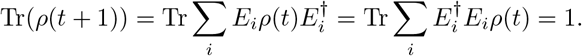

Since above equation is true for all *ρ* then

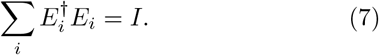

From the definition of Kraus operators in Eq. (6), a step of the Bcd walker is given by

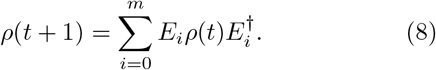

Fro *t* steps of the evolution, one can then write

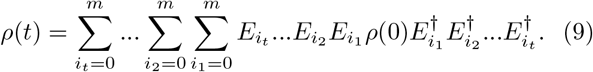

The coin operator, the translation operator and the environment effects are all embedded in *E*_*i*_. Our task is then reduced to finding the appropriate Kraus operators for our system defined in Eq. (6) and then use Eq. (9) in order to obtain the final state *ρ*(*t*). The Kraus operators naturally take the influence of the environment on the system into account. In our case this can be shown as follows. Le *R*_*i*_ is a reaction operator that acts on the system with the corresponding probability *p*_*i*_. Furthermore, let us assume that we have *r* such operators *R*_*i*_, *i* = 1, …, *r*, where each of them acts on the system with the probability *p*_*i*_. Now, if the environment is in the state |*e*_*i*_⟩ then the operator *R*_*i*_ acts on the system. Therefore, one can imagine the *r*-dimensional Hilbert space for the environment, spanned by 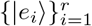, and the following initial state for the environment

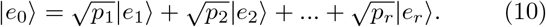

Clearly, the probability of finding the environment in the state | *e*_*i*_ ⟩ is *p*_*i*_. Therefore one can write the unitary transformation of the whole system (system+environment) as

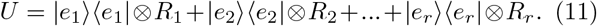

So the Kraus operators will be given by

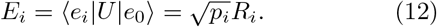

and Eq. (8) gives the density matrix after the first step accordingly as

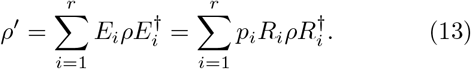

Equation (13) gives *ρ* as a mixture of different evolutions with the corresponding probabilities *p*_*i*_ as expected. Translating this to our setting of the noise model given in Fig. (2) we can define the initial state of the environment as follows

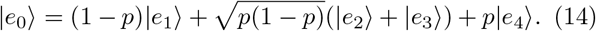

Since *E*_*i*_ are the operators that act on the system (coin+position) Hilbert space, one can thus write the general form of *E*_*i*_ as follows

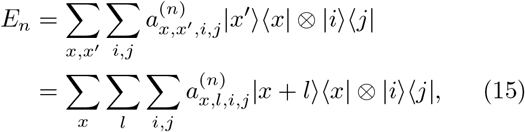

where *x, l* = −∞,…, *∞* and *i, j* ={|™⟩, |+⟩}.Fourier transformation

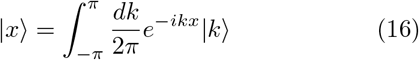

turns Eq. (15) into

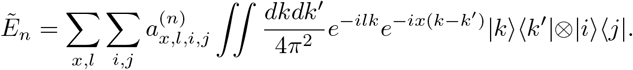

After invoking the homogeneity condition so that the coefficients 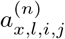 takes the form are independent of *x*, above equation

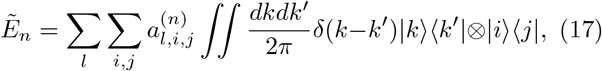

where we have made use of the following orthonormalzation relation

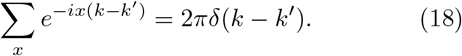

By integrating out the terms involving *k*^*′*^ and changing the order of integration and summation one gets

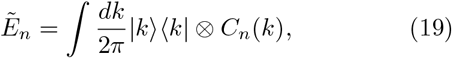

where

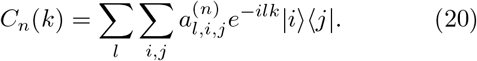

One may, in a similar manner, write the general form of *ρ* in the *k* space as

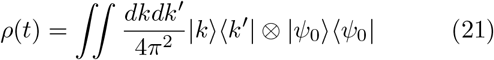

so that Eq. (8) becomes

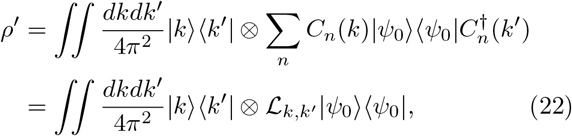

where L_*k,k*_*′* is a superoperator defined as

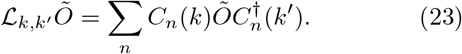

Thus after *t* steps, the state and the probability of finding our Bcd walker at position *x* are, respectively, given by

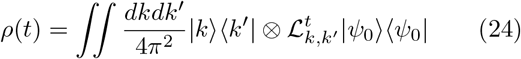

and

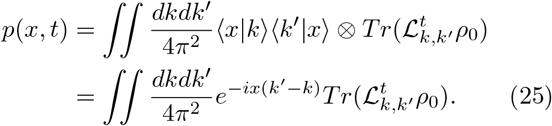

From the completeness relation, Eq. (7), one can thus deduce that

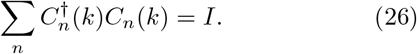

From the above condition it follows that the superoperators L_*k,k*_ are trace preserving, i.e.,

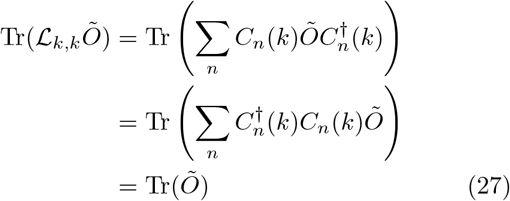

and therefore

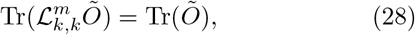

which is true for any arbitrary operator *Õ*.

Now the *m*th moment of the probability distribution *p*(*x, t*) is defined by

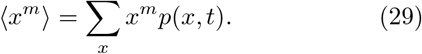

Inserting Eq. (25) in Eq. (29) we get

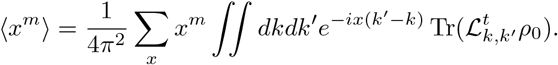

Using the orthonormalization relation (18), one gets for the first and second moments

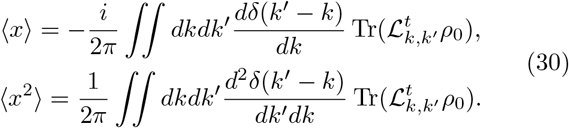

In order to carry out the above integrations one needs the following equations

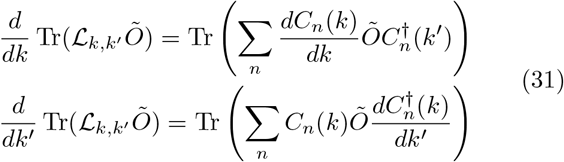

where according to Eq. (20)

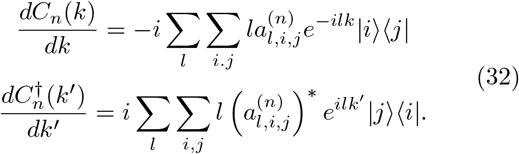

Since the superoperator _*k,k*_*′* acts on a positive and Hermitian density matrix, one can write Eq. (31) as follows

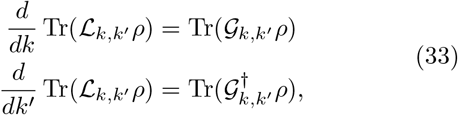

where

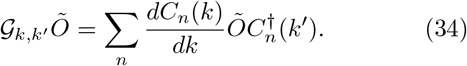

Finally by carrying out the integrations of Eq. (30), we arrive at the following expressions for the first and second moments

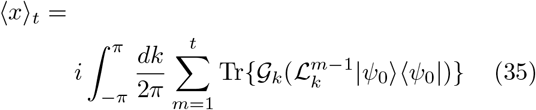

and

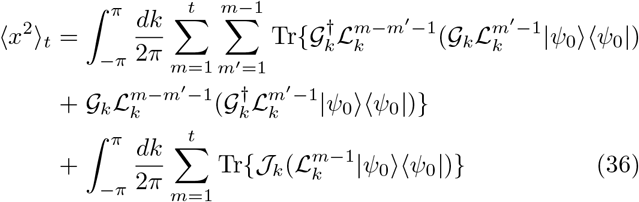

where we have defined

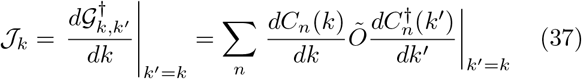

and used L_*k*_ ≡ L_*k,k*_*′* and 𝒢_*k*_ ≡ 𝒢_*k,k*_*′* to simplify writing. Now, following Ref. [45], we use the affine map approach in order to find the above superoperators and determine the value of Bcd diffusivity as follows. Since the operator L_*k*_ is linear one can represent it as a matrix acting on the space of two-by-two operators. First, any twoby-two matrix can be represented by a four-dimensional column vector as

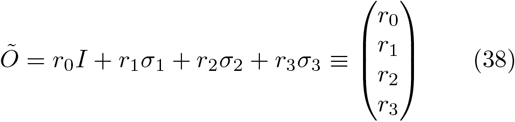

where we have defined *σ*_0_ = *I* and *σ*_*i*_ (*i* = 1, 2, 3) are the usual Pauli matrices. Now, in order to find the above superoperators, we let them operate on an arbitrary twoby-two matrix Õ as follows

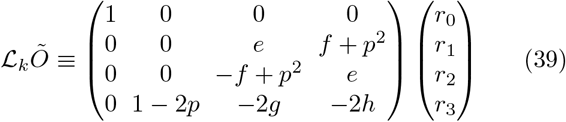

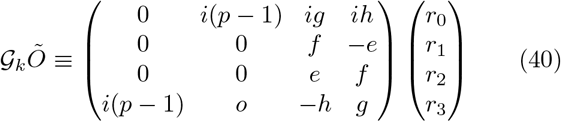

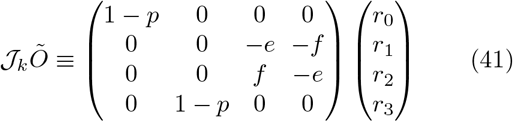

and

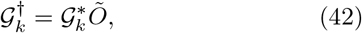

where the last equation is obtained, simply, from the Hermiticity of the Pauli matrices. Here *e, f, g*, and *h* are functions of *p* and *k*, defined by

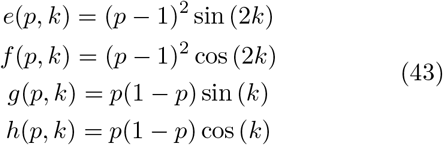

With the above representation in hand, one can calculate the moments given by Eq. (30). It turns out that, in the long time limit, the first moment is independent of time. Our interest here is in finding the diffusion coefficient with the below definition

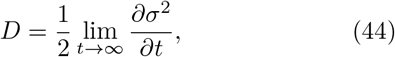

where *σ*^2^ = ⟨*x*^2^⟩ −⟨*x*⟩ ^2^. Since the time independent term does not contribute to the diffusion coefficient *D*, thus we shall focus on finding the second moment in Eq. (30). A somewhat detailed but straightforward calculation shows that the second moment is given by

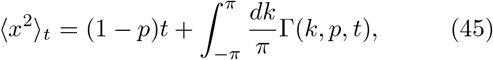

where

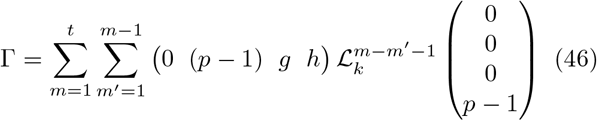

Substituting the result in Eq. (45) into the definition of diffusion coefficient in Eq. (44) and simplifying we have finally

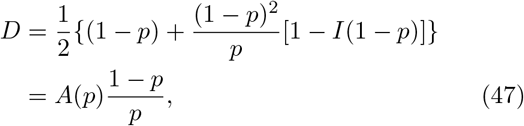

where

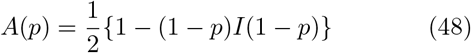

and

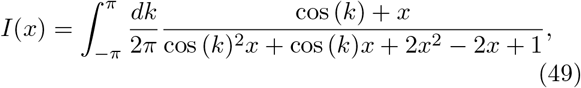

where the value of the coefficient *A→* 0.5 as *p →*1. Remarkably, Crick, using his source-sink model,

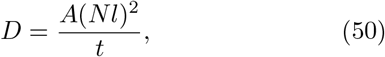

has proposed a general value of *A* = 0.5 for the biochemically more realistic models, where *t* is the time, *N* is the number of cells between the source and sink, and *l* is the length of each cell [16]. We shall now show that the higher values of local diffusivity for *Bcd* allowed by Eq. (47) (see Fig. 3) can reduce the search times for binding to the promoter sites on the hunchback *hb* loci by allowing higher sliding and association rates along the DNA leading to further enhancement in the precision of gene expression boundaries.

**FIG. 3.**
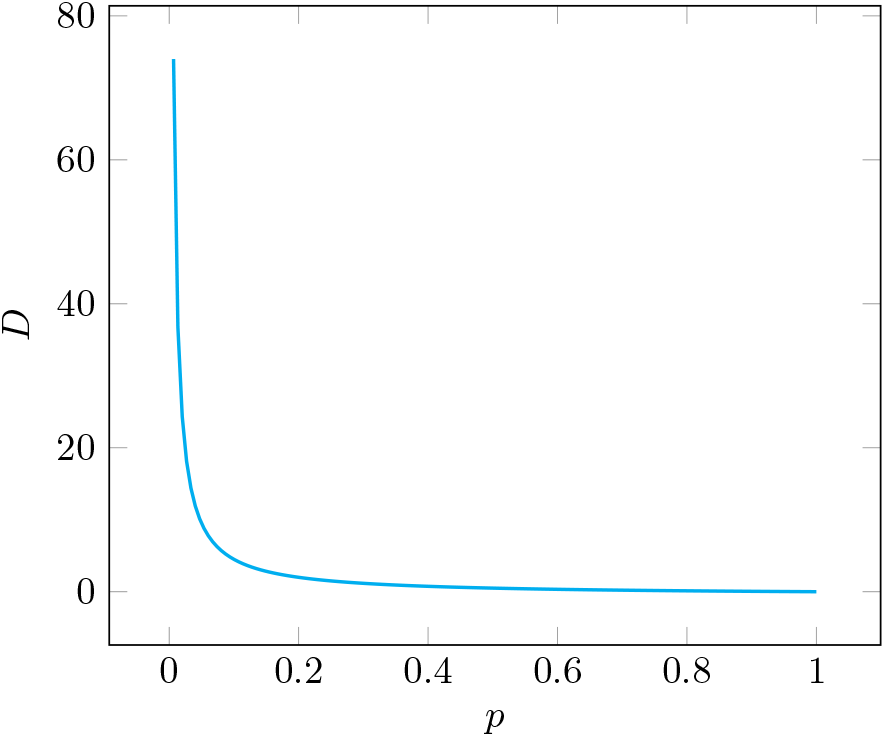
Plot of Eq. (47). The magnitude of *D* is large for very small *p* and we have taken *A* = 0.5.

*Bcd* is an example of a morphogen that directly acts on the process of transcription since it is also a transcription factor [46]. It is a maternally expressed protein involved in body patterning along the anterior-posterior axis in the early Drosophila embryo [14]. *Bcd* has at least 66 target genes including hunchback (*hb*) [47]. It elicits a remarkably fast and precise transcriptional response. Gene expression in response to the *Bcd* nuclear import gets established in only a couple of minutes [48]. This fact points to a very efficient search mechanism to locate the target genes. At the same time, small variations in concentration along the embryo anterior-posterior axis are enough to turn on or off the transcription of its target genes, leading to well-defined expression domains [49]. The *Bcd* search times for its binding sites on the *hb* promoter gene that are usually employed by the classical models of *Bcd* gradient interpretation [50] are estimated by assuming a 3*D* search process inside the nucleus space [51]. The trade-off faced by such approaches is that the reduction in the binding times of the transcription factor to the promoter sites demands diffusion coefficients which are an order of magnitude larger than are possible if the motion of Bcd molecules takes place only through a classical random walk based mechanism [52]. However, a model based on a quantum walk has no such limitation as the diffusivity of the system can easily attain values much larger than are possible classically depending on the noise level at a given point in time (Fig. 3).

The large values of diffusivity possible in a quantum walk model can also avoid the paradox of faster-than-diffusion association rates without any need for the transcription factor to constantly alternate between 1*D* and 3*D* diffusion-based search processes [53]. The measured association rate *κ* ∼ 10^10^*M* ^*−*1^*s*^*−*1^ for the LacI repressor with the promoter sites of DNA in bacteria was found to be much larger than that allowed by the classical laws based on a 3*D* diffusive motion [54]. For instance, the well-known Debye-Smoluchowski equation for a bimolecular association process

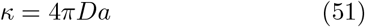

predicts a maximal rate of ∼ 10^8^*M* ^*−*1^*s*^*−*1^, where *a* denotes the receptor size [53]. In order to resolve this paradox, the hypothesis of rapid search by the protein sliding along the DNA in a 1*D* diffusion process and alternating between rounds of 3*D* diffusion in the bulk was proposed [55]. However, in a model based on quantum diffusion rather than a classical one the developing fly embryo can avoid the fate of very low association rates by exploiting quantum effects. Substituting the expression for *D* from Eq. (47) into Eq. (51) gives

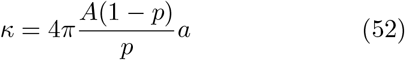

Putting in the value of receptor size *a*∼ 3*nm* from Ref. [56] and the numerical value of noise *p* comparable in magnitude to the observed *Bcd* degradation rate (*k* ∼ 10^*−*4^*s*^*−*1^) [57] in above equation shows that *κ*∼ 10^10^*M* ^*−*1^*s*^*−*1^. This is the least value that the *Bcd* association rate can have in the organism, considering that the eukaryotic protein-DNA recognition, unlike the case in bacteria, is already complicated by chromatin packing of the DNA and the multisubunit structure of the transcription factors [53], and this association rate is comparable to that observed for bacteria [54].

In conclusion, we have used an open quantum system approach to analyse the dynamics of Bcd gradient formation and shown that exactly the same dynamics are obtained through this more rigorous method. Using the thus obtained expression for the diffusion coefficient, we have shown that the large values of diffusivity allowed by quantum mechanics can avoid the paradox of faster-than-diffusion association rates without any need for the transcription factor to constantly alternate between 1*D* and 3*D* diffusion-based search processes. Since many transcription factors are known to share a common search strategy for target gene regulatory regions, the mechanism we propose here may have a wide range of applicability in biological processes.

I would like to thank Carl Trindle for financial support.

